# Trait and Growth Responses to Sequential Environmental Change linked to Sensitivity in *Synechococcus* populations

**DOI:** 10.1101/2025.06.11.659080

**Authors:** Arunima Sikder, Colin T Kremer, Frederik De Laender

## Abstract

1. Environmental change often occurs as a sequence of stressors rather than as isolated events. While the individual and combined effects of multiple stressors are well studied, the ecological consequences of sequential environmental change remain poorly understood. Such sequences, for example, a marine heatwave followed by seasonal herbicide runoff, are increasingly common under global change.
2. We investigated how legacy effects (the imprint of past environmental conditions) influence subsequent population performance and functional traits, and how these effects are mediated by strain-specific sensitivities.
3. Using a fully factorial design, we exposed six strains of the globally abundant pico-phytoplankton *Synechococcus* sp. to three environmental conditions: warming, herbicide exposure, and a control, under both chronic (same condition across time) and sequential (different conditions across time) regimes. We measured population performance (per-capita growth rate, maximum density) and key functional traits (cell size, chlorophyll content).
4. Population responses diverged significantly between chronic and sequential exposures, revealing a strong legacy effect. Trait changes were often decoupled from growth metrics, suggesting independent response axes. Strain identity and its interaction with past conditions explained substantial variation in both growth and trait responses.
5. We identify and conceptualise four distinct mechanisms of legacy effects during sequential change: overcompensation, amplification, constraint and depression, each linked to strain-specific responses. Consequently, incorporating legacy effects into predictions of biodiversity dynamics and ecosystem function under global change is therefore both feasible and essential.

## 1. Introduction

Environmental drivers of population growth vary through space and time, and global change often amplifies this variability (Shibley et al., 2002). These shifts of intensity, frequency, and duration of environmental drivers can significantly influence population growth and functioning (Oliver et al., 2021; Xu et al., 2022). Population responses to individual changes in key drivers, such as temperature (Eppley 1972; Thomas et al. 2012; Bernhardt et al., 2018) or nutrient availability (Droop, 1973; Borer et al., 2017; Hillebrand et al., 2014), are well characterised. Additionally, the interactive effects of co-occurring drivers have been explored through meta-analyses (Orr et al., 2024; 2020) or by experimentally manipulating the number and similarity of drivers (Rilling et al., 2019; Holmes et al., 2021). However, natural systems more often experience sequential perturbations rather than isolated or simultaneous changes (Nguyen et al. 2020). For example, in marine environments, heatwaves create alternating periods of warming and cooling (Wolf et al., 2024), which may be followed by seasonal influx of pollutants (Jaikaew et al., 2015). Such sequences create complex spatiotemporal mosaics of environmental conditions, to which population responses remain largely unknown. A major factor in determining population response to sequential exposure is the legacy of past environments (Jackson et al., 2021). Legacy effects describe how previous biotic or abiotic conditions influence current population performance (Monger et al., 2015). For example, legacy effects have been detected in lichen communities persisting for up to 16.5 years (Johansson et al., 2018), and in marine biofilms where temperature fluctuation history enhances stability during subsequent extremes (Rindi et al., 2025). Despite these insights, the mechanisms by which legacy effects shape population performance remain underexplored.

Understanding legacy effects is challenging because they are often mediated by functional trait plasticity or acclimation, including delayed shifts in trait means that enable populations to adjust to changing conditions and influence responses to subsequent environments (Coates et al., 2025). These shifts can be gradual or synchronous, reversible or irreversible, depending on the nature and timing of environmental change (Schulte et al., 2011; Kremer et al., 2018). When environmental conditions shift sequentially and differ in nature, trait changes may or may not be fully optimised, thereby affecting population performance. Disentangling legacy-driven plasticity therefore requires comparing trait changes under varying versus constant conditions, while simultaneously tracking trait dynamics and population growth. In this study, we compare population performance between chronic exposures, where past and subsequent environments remain the same (XX), and sequential exposures, where environments change (XY). While previous work has linked functional trait shifts to demographic outcomes (Fey et al., 2021; Holmes et al., 2024; Bruin et al., 2023; Wieczynski et al., 2021), it remains unclear how prior environmental conditions influence subsequent trait expression and growth under such sequential exposures.

We hypothesise that the strength and direction of legacy effects during sequential exposure are shaped by an organism’s sensitivity to environmental change (Schuetz et al., 2019). Here, we define sensitivity as the deviation in growth or trait expression when populations are chronically exposed to a non-reference environment, relative to a control. Sensitivity may be either positive or negative, depending on whether performance or trait values are amplified or depressed in the altered environment (Fig. 1), and can vary substantially among strains or populations.

**Fig 1:**
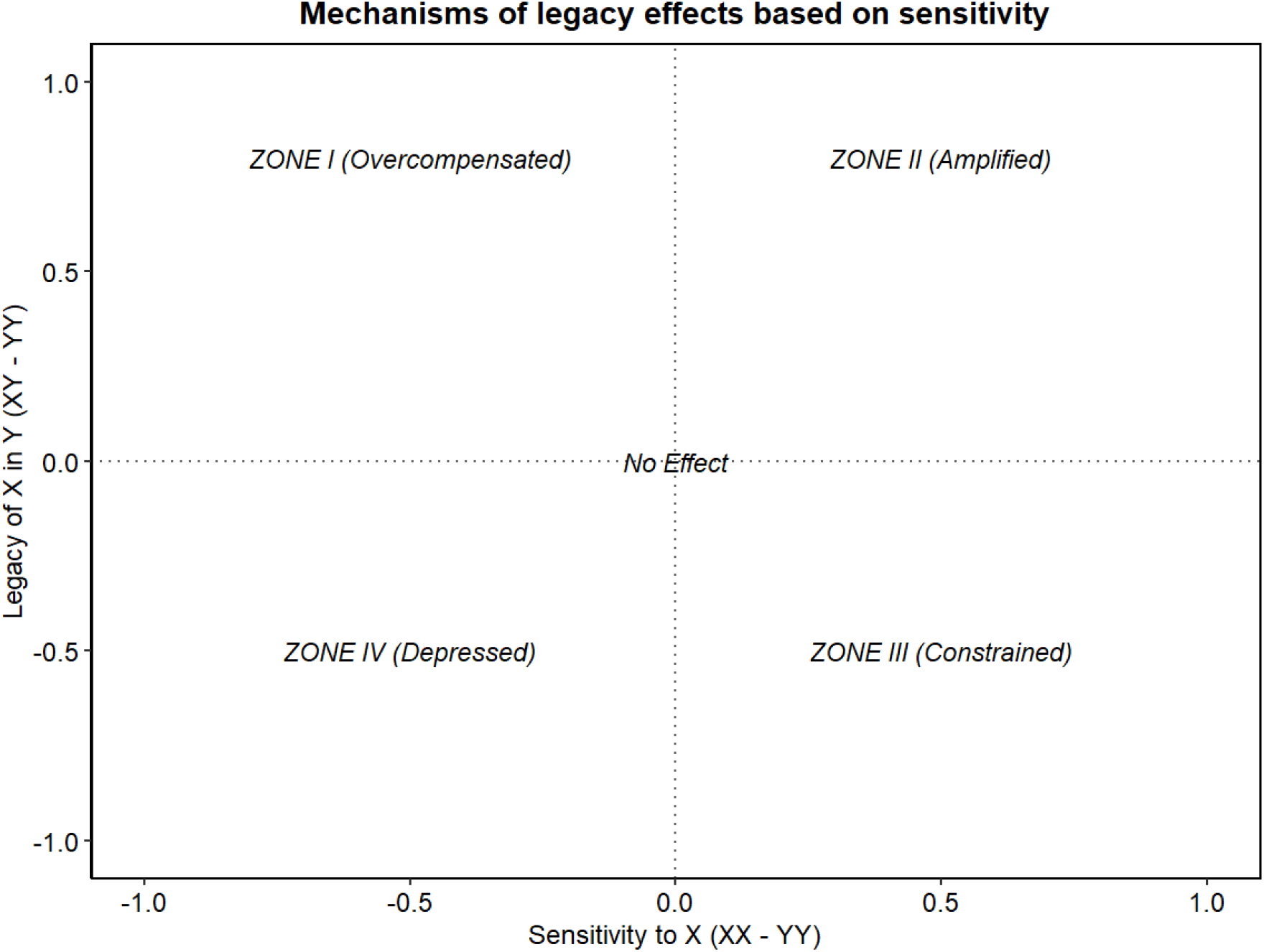
Conceptual plot illustrating links between sensitivity (x-axis) and legacy effects (y-axis) for population growth and traits during sequential change. Sensitivity (x-axis) – difference in growth and traits under chronic X compared to under chronic Y (XX-YY). Legacy effect (y-axis) – influence of prior exposure to X on growth and traits in subsequent Y. **Zone I**: **Overcompensation** – sensitivity to X is negative (reduced growth and traits under chronic X than Y) but legacy of X is positive. **Zone II**: **Amplification** – sensitivity to X is positive and legacy of X is also positive thereby boosting performance under Y. **Zone III**: **Constrained** – positive sensitivity is dampened by a negative legacy effect. **Zone IV: Depressed** – both sensitivity and legacy effect are negative leading to depressed growth and trait expression. **No effect** – rapid plasticity may minimize differences between XY and YY, leading to negligible legacy effects.

The goal of this study is to evaluate legacy effects: how does the past environment in a sequence affect population growth and traits in the subsequent environment, and are these effects related to strain-sensitivity? Studying a globally relevant phytoplankton genus, we investigated responses under three environmental conditions: a control (C; reference conditions), warming (T; + 2°C), and herbicide pollution (P; atrazine). We quantified population responses using two growth metrics, density and per-capita growth rate (PCGR), and two functional traits linked to ecosystem processes, cell size (metabolic rate and biomass, Hinners et al., 2024; Marañón, 2015) and chlorophyll content (light harvesting and productivity, Gui & Sun, 2024). We use a fully-factorial experimental design crossing all past and subsequent environments (Fig 2).

**Fig 2:**
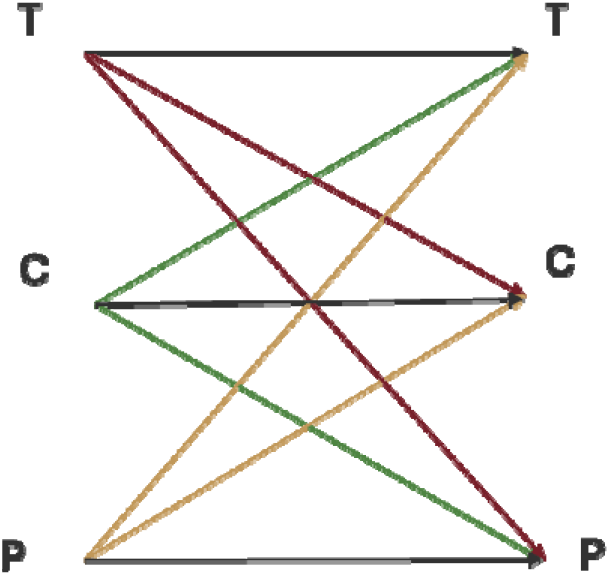
Schematic of experimental design for all environmental exposures investigated in thi study using control (C), pollution (P), warming (T). Chronic exposures (same past and subsequent environment): C→C, P→P, T→T (black arrows). Sequential exposure (mismatched past and subsequent environments): C→P, C→T (green arrows), P→C, P→T (yellow arrows), T→C, T→P (red arrows). Arrow direction indicates sequence order (past → subsequent environment).

Through this experimental design, we tested two hypotheses. H1: if past environmental conditions influence population growth in subsequent environments, it will also induce changes in functional traits, as traits mediate organismal responses to environmental variability. H2: the legacy of past environments on population growth and traits will be greater in strains that are inherently more sensitive to that specific environment.

### Model Organism

The focal genus of this study, *Synechococcus* sp., is a globally dominant marine phytoplankton belonging to the taxa cyanobacteria. It contributes substantially to marine primary production and nutrient cycling (Wang et al., 2022; Chen et al., 2022). *Synechococcus* sp. is an ideal model for investigating population responses to environmental shifts due to its short generation times and global dominance (Allaf et al., 2022). The six strains in this study span different habitats of origin and phycoerythrin-to-phycocyanin pigment ratios, reflecting adaptations to varying light regimes at different ocean depths (Grébert et al., 2022; Table 1). For this study, we expect such adaptations to result in varying degrees of sensitivities to the environmental drivers concerned. In marine ecosystems, both pollution and warming are significant environmental drivers, which cause photosynthetic inhibition (Wang et al., 2024; Chalifour & Juneau, 2011) and alter phytoplankton growth rates, diversity, and community structure (Henson et al., 2021). Consequently, understanding phytoplankton responses to sequential exposures of these drivers is critical for predicting impacts on marine ecosystem function.

**Table 1:**
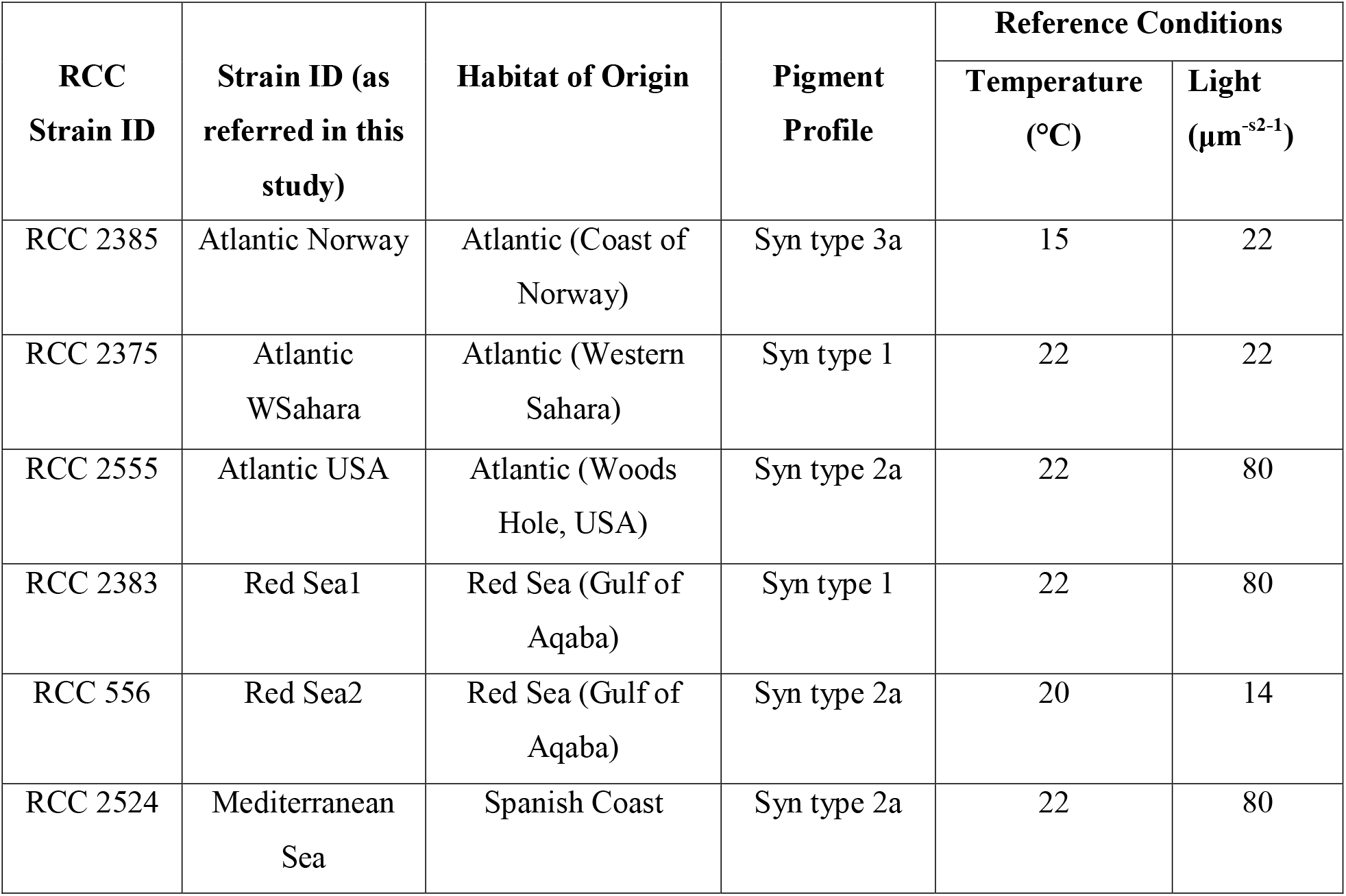
Origin and laboratory culture conditions of the focal strains of *Synechococcus* sp. Information derived from Roscoff Culture Collection website.

| RCC<br>Strain ID | Strain ID (as<br>referred in this<br>study) | Habitat of Origin | Pigment<br>Profile | Reference Conditions |  |
| --- | --- | --- | --- | --- | --- |
| | | | | Temperature<br>(°C) | Light<br>($\mu\text{m}^{-2}\cdot\text{s}^{-1}$) |
| RCC 2385 | Atlantic Norway | Atlantic (Coast of<br>Norway) | Syn type 3a | 15 | 22 |
| RCC 2375 | Atlantic<br>WSahara | Atlantic (Western<br>Sahara) | Syn type 1 | 22 | 22 |
| RCC 2555 | Atlantic USA | Atlantic (Woods<br>Hole, USA) | Syn type 2a | 22 | 80 |
| RCC 2383 | Red Sea1 | Red Sea (Gulf of<br>Aqaba) | Syn type 1 | 22 | 80 |
| RCC 556 | Red Sea2 | Red Sea (Gulf of<br>Aqaba) | Syn type 2a | 20 | 14 |
| RCC 2524 | Mediterranean<br>Sea | Spanish Coast | Syn type 2a | 22 | 80 |

## 2. Materials and Methods

### 2.1 Experimental design and procedure

All strains of *Synechococcus* sp. (Table 1) were obtained from the Roscoff Culture Collection (Brittany, France) and inoculated under aseptic conditions into 50 mL Erlenmeyer flasks containing PCR-S11 Red Sea medium following Rippka et al. (2000). Stock cultures were maintained in thermostatic cabinets (Lovibond) fitted with platform shakers (Fisher Scientific) at 150 rpm, under a 12 h:12 h light: dark cycle at reference conditions for each strain (Table1).

The experiment comprised two phases: phase 1 involved exposure to the first environment (hereafter past environment), followed by phase 2 exposure to the subsequent environment. For chronic exposures, environments remained constant across both phases, whereas for sequential exposures, environments changed between phases (Fig. 2). We initiated the experiment in microcosms using stock cultures of the six strains. Six-well culture plates (Corning Incorporated Costar) served as microcosms with a total volume of 6mL per well. Inoculation for phase 1 was done by adding 2 mL of sample from stock cultures (30,000 cells mLD^1^) and 4mL of appropriate media (PCRS11 or polluted) as per experimental setup. Phase 1 lasted seven days to capture a complete growth curve for *Synechococcus* strains (mean generation time ≈ 24 h). Before starting phase 2, all cultures within each exposure group and strain were diluted to a common density to ensure consistent initial population sizes across strains. Thereafter, replicates of each strain-culture were reinoculated into three different environmental conditions, resulting in 162 sub-cultures. These sub-cultures were allowed to grow for 10 days, which was the time it took to reach carrying capacity in most treatments. Plates were sampled every 24 h for 17 days: at each sampling, 200 µL from each well of a microcosm was withdrawn aseptically into a 96-well plate (Corning Incorporated Costar) and replaced with 200 µL of pre-equilibrated media to maintain constant volume and pollutant concentration.

Population density, cell size, and chlorophyll content were quantified by flow cytometry (Guava easyCyte HT-7) using a 488 nm excitation laser, Forward Scatter (FSC detector), and RED-B channel (610/20 nm emission) respectively. Debris and doublets were excluded via gating in GuavaSoft 4.5 based on FSC-H vs. FSC-A thresholds. Raw .fcs files were imported into R (v 4.4.2; R Core Team 2023) via the flowCore (v2.14.2, Elis B et al., 2024) and cyanoFilter (v1.10.0, Olusoji et al., 2021) packages for post-hoc quality control (debris gating, doublet removal).

### 2.2 Data treatment and Statistical analysis

All statistical analyses were conducted in R Statistical Software (v4.4.2; R Core Team 2023). From the phase 2 data, we extracted four response variables per replicate: population density, perDcapita growth rate (PCGR, dayD^1^), cell size and chlorophyll content. For all variables except PCGR, log-transformed values were modelled as smooth functions of time using Generalized Additive Models (GAMs) with the “mgcv” package (v1.9-3; Wood 2011). These models treat the mean logarithm of each response variable as a smooth function (f) of time (t).

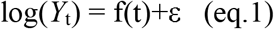

where Y_t_ is the response at time t, f is a thin-plate regression spline with basis dimension k =7, and smoothing penalties estimated by restricted maximum likelihood (REML) to balance flexibility with parsimony (Wood 2011). Using predicted values from the fitted GAM for density, we calculated per-capita growth rate (PCGR) over each time interval for each replicate as

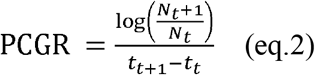

Here, *N*_*t*_ is the fitted population density at time *t*_*t*_, *N* _*t*+l_ is the fitted population density at the next time step, and *t*_*t*+l_ -*t*_*t*_ is the time interval between observations. Data manipulation and summary statistics were performed using the “tidyverse” package (v2.0.0; Wickham et al. 2019). Data treatment also included Principal Component Analysis (PCA) of centred and scaled response variables to assess sample clustering. Biplots were generated via the “ggfortify” (Tang et al., 206) package with loading vectors overlaid to indicate variable contributions.

Statistical analyses were conducted to assess variation in population responses between chronic and sequential exposures. For each replicate, trait values were averaged over the 10-day Phase 2 period, while growth metrics (PCGR and density) were summarized as maximum values. Analyses were performed separately for chronic and sequential exposures. Mixed-effects ANOVAs was conducted using the “afex” package (v1.4-1; Singmann 2024), with replicate as the subject ID, exposure type as a within-subject factor, strain as a between-subject factor, and including the strain × exposure type interaction term. Each response variable was analysed independently using type III sums of squares. Results are presented as type III ANOVA tables with Generalized Eta-Squared (ges) reported for all effects. Post-hoc estimated marginal means were calculated with the “emmeans” package (v1.11.1; Lenth 2025), applying Tukey-adjusted pairwise contrasts (α = 0.05) between exposure types for each strain. Significance was denoted as p < 0.01 (**), p < 0.05 (*), and p < 0.10 (.). To quantify strain sensitivities, chronic exposures (PP, TT) were compared to control (CC). Legacy effects were quantified by contrasting sequential exposures with their corresponding chronic references (e.g. PC vs. CC), thereby testing hypothesis H1. For hypothesis H2, we assessed whether the magnitude and direction of legacy effects covaries with a strain’s sensitivity to that same environment for all sequential exposures. Results were interpreted within the framework outline in Fig. 1, which illustrates how sensitivity and legacy effects may manifest in *Synechococcus* sp. populations exposed to sequential environmental regimes.

## 3. Results

### 3.1. Population and trait expression reveals strain-specific sensitivities under chronic exposure

Chronic exposures to warming and pollution significantly affected trait and growth metrics (Table 4), with strong effects of strain identity (F(5, 10) = 55.0, p < 0.001), exposure type (F(2, 20) ≈ 111, p = 0.002 to < 0.001), and their interaction (strain × exposure: F(9.88, 19.8) = 54.0, p < 0.001).

Both maximum density and per capita growth rate (PCGR) declined consistently under chronic pollution, while responses to chronic warming were more variable across strains (Fig. 3). Cell size increased moderately under warming (+ 0.5, *p* = 0.002), consistent with warming-induced physiological shifts, whereas chlorophyll content showed strain-specific increases or decrease (Fig. 3).

**Figure 3.**
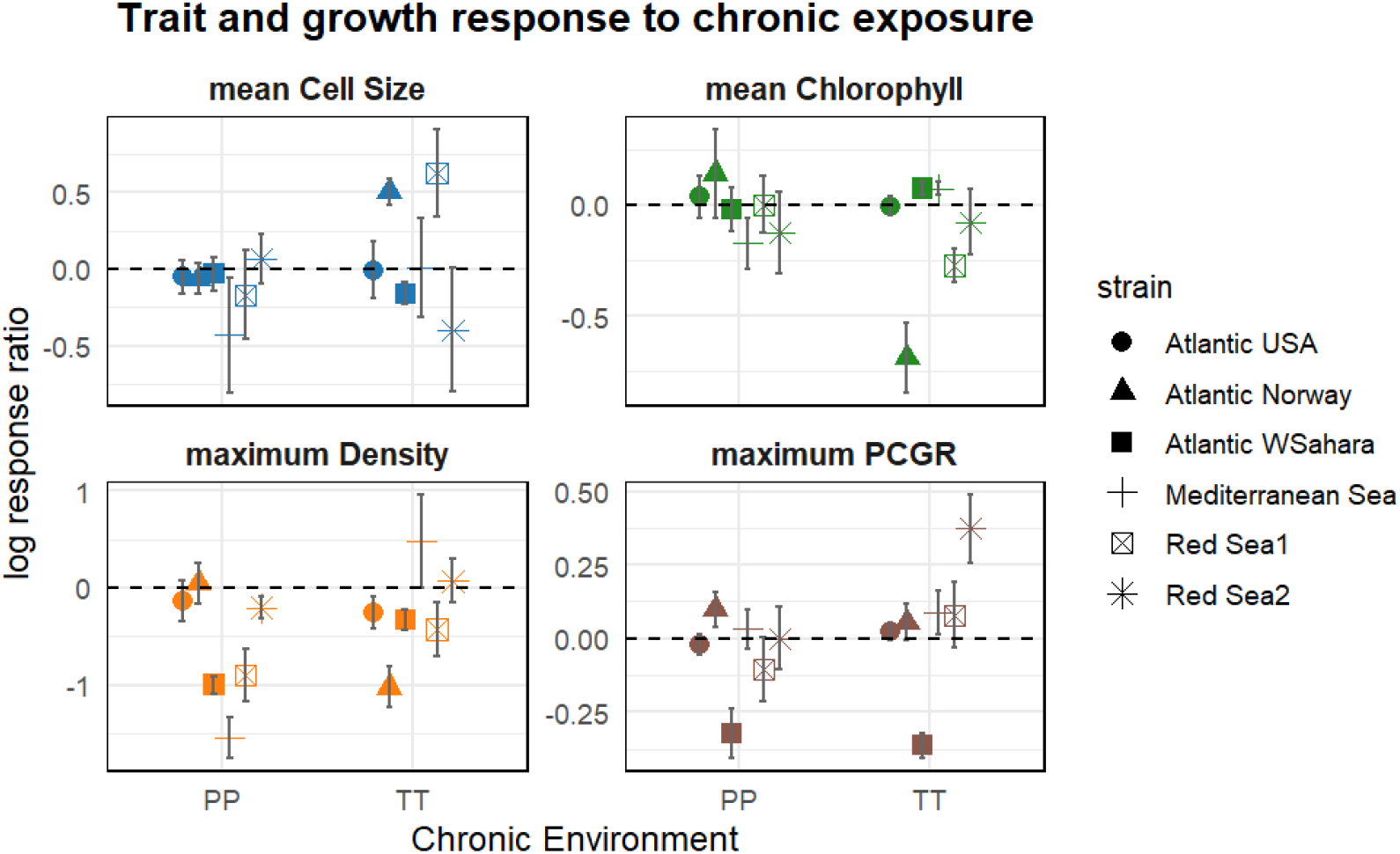
Strain-specific responses to chronic pollution and warming in six strains of *Synechococcus* sp., shown as the mean log-response ratios (n=3, ± 95% confidence intervals) relative to the control. Each panel displays strain-level responses for four variables: cell size, chlorophyll content, population density, and per capita growth rate (PCGR). Log-response ratios were calculated as the natural log difference from control conditions (e.g., log(TT) – log(CC) or log(PP) – log(CC)). A log-response ratio of zero (dashed horizontal line) indicates no sensitivity. Shapes represent strain identity; colours indicate response variables (cell size, chlorophyll content, population density, and PCGR).

Strains originating from the Atlantic exhibited higher sensitivity to chronic warming (Fig. 3), while those from warmer habitats were more sensitive to chronic pollution. These patterns suggest that the relative impact of different chronic environments is strain-dependent and may reflect local adaptations.

Interestingly, we observe divergence between maximum density and PCGR: some strains exhibited high maximum PCGR but failed to reach high maximum densities, likely due to early resource depletion or stress sensitivity. Conversely, others maintained slower but more sustained growth, achieving high maximum final densities despite lower PCGR. Moreover, trait shifts did not always align with growth outcomes, indicating decoupling of growth and trait strategies among *Synechococcus* strains. This divergence is corroborated by our PCA analysis (Supplementary Materials Fig. 1), which shows that the four response variables loaded on separate axes, further supporting functional decoupling. Together, these results reveal that (i) sensitivity is both strain-specific and dependent on the type of environmental driver, and (ii) cell size, pigment content and growth metrics exhibit divergent response strategies under chronic exposure within this experimental timeframe.

### 3.2. Legacy effects on population growth and traits during sequential exposure

Sequential exposures revealed pronounced legacy effects on both growth metrics and functional traits, with past environmental conditions significantly shaping population performance under subsequent exposures (Supplementary Materials, Table 2). Under subsequent control conditions (C), past environments significantly affected density (*F*(1.73, 19) = 16.7, *p* < 0.001), PCGR (*F*(1.73, 19) = 16.7, *p* < 0.001), and chlorophyll content (*F*(1.64, 18) = 19.3, *p* < 0.001), demonstrating persistent legacy effects even when populations returned to reference conditions. Similarly, under subsequent pollution conditions (P), past environmental exposures significantly influenced density (*F*(1.83, 18.3) = 21.3, *p* < 0.001), cell size (*F*(1.14, 11.4) = 22.7, *p* < 0.001), and chlorophyll content (*F*(1.47, 14.8) = 8.85, *p* = 0.005). For subsequent warming (T), past environments had significant, though comparatively smaller, effects on cell size (*F*(1.16, 10.4) = 5.03, *p* = 0.044), PCGR (*F*(1.59, 14.3) = 16.9, *p* < 0.001), and chlorophyll (*F*(1.87, 16.8) = 15.1, *p* < 0.001).

**Table 2:** Environmental conditions and their descriptions for this study.

| Environmental Condition | Description |
| --- | --- |
| Control (C) | Reference temperature, light and growth media (PCRS11). |
| Pollution (P) | Reference temperature, light and growth media (PCRS11) with atrazine (0.029 mgL<sup>-1</sup>). |
| Warming (T) | Reference light and growth media with 2°C increase from reference temperatures (as in Table 1). |

Additionally, strong and consistent strain × past environment interactions across most responses (all p < 0.01) indicate that these legacy effects are modulated by strain-specific adaptations. In line with our hypothesis H1, these results demonstrate that legacy effects significantly influence growth rates and trait expression in a manner that is both strain and environment-specific. Together, these findings robustly support the existence of legacy effects whereby past environmental exposures significantly shape subsequent growth and functional trait expression.

### 3.3 Sensitivity explains mechanisms of legacy effect on growth and traits

In line with hypothesis H2, strains that were sensitive under chronic exposures also exhibited pronounced legacy effects during sequential exposures.

The magnitude and direction of legacy effects across sequential exposures were closely associated with strain-specific sensitivity to each environment, revealing distinct patterns for growth metrics and functional traits (Fig. 5). Whenever there was a non-zero difference in traits and growth due to chronic exposure (x axis, sensitivity), corresponding changes due to legacy effects were observed (y axis Δ ≠ 0). Across all sequences, trait and growth metrics predominantly clustered in zones II and IV. This pattern indicates that high environmental sensitivity often resulted in amplification (zone II) or depression (zone IV). Less frequently, but notably, overcompensation (zone I) was observed due to a legacy of pollution in PT and PC sequences, and warming in the TP sequence. Additionally, constrained responses (zone III) were linked to legacy of control conditions in subsequent pollution and warming exposures (i.e., CP and CT sequences).

**Figure 4:**
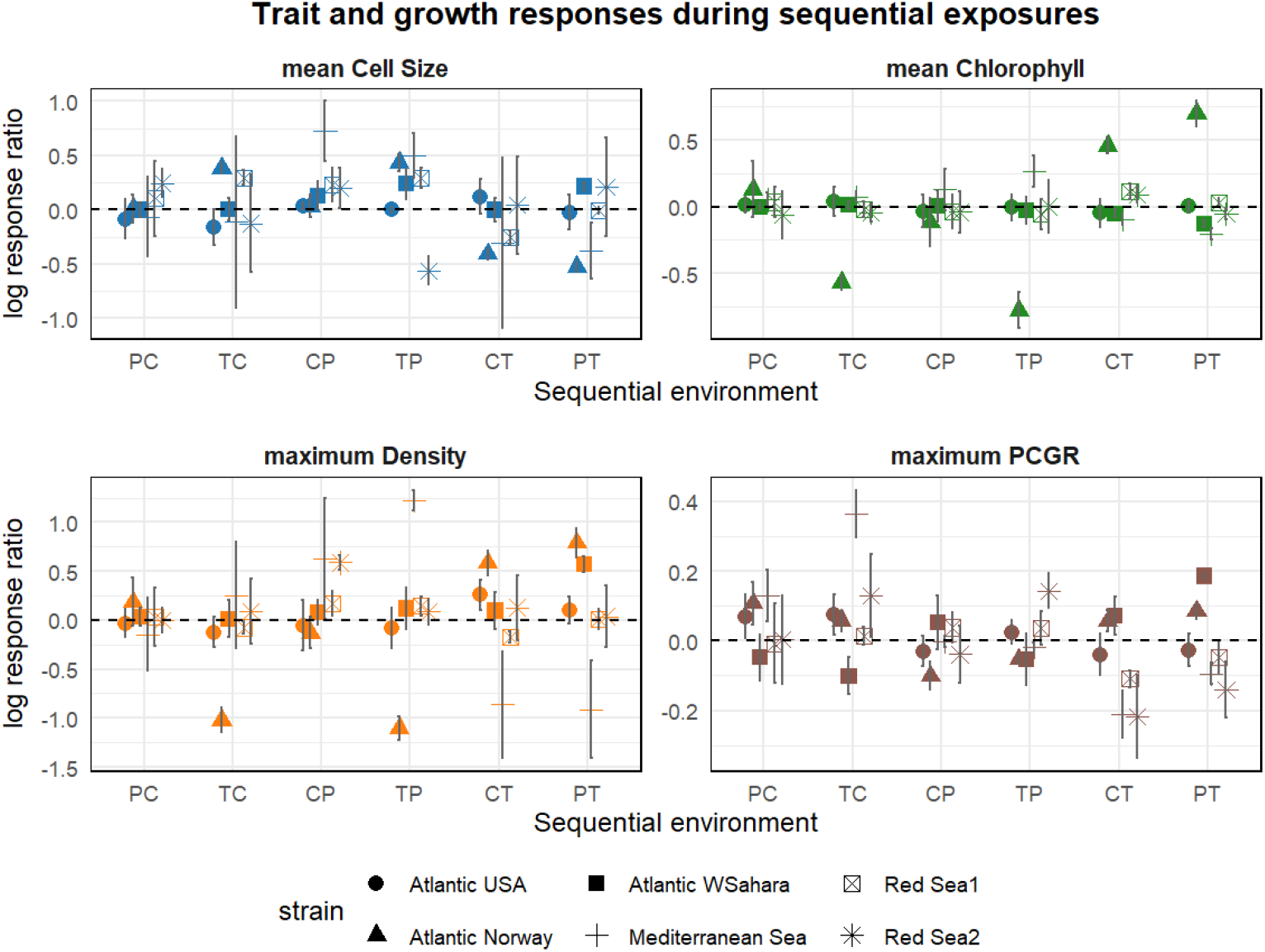
Legacy effects of past environments during sequential exposure on trait and growth metrics in six strains of *Synechococcus* sp. Environmental conditions (X and Y) include pollution, warming, and control, resulting in six sequential environments (X→Y, shown on the x-axis). Each panel displays mean log-response ratios (n = 3, ± 95% confidence intervals) comparing sequential exposures (XY) to chronic exposures (YY). A log-response ratio of zero (dashed horizontal line) indicates no legacy effect. Shapes represent strain identity; colours indicate response variables (cell size, chlorophyll content, population density, and PCGR). Post-hoc contrasts for the associated ANOVA test is available in Supplementary Materials Table 3.

**Fig. 5.**
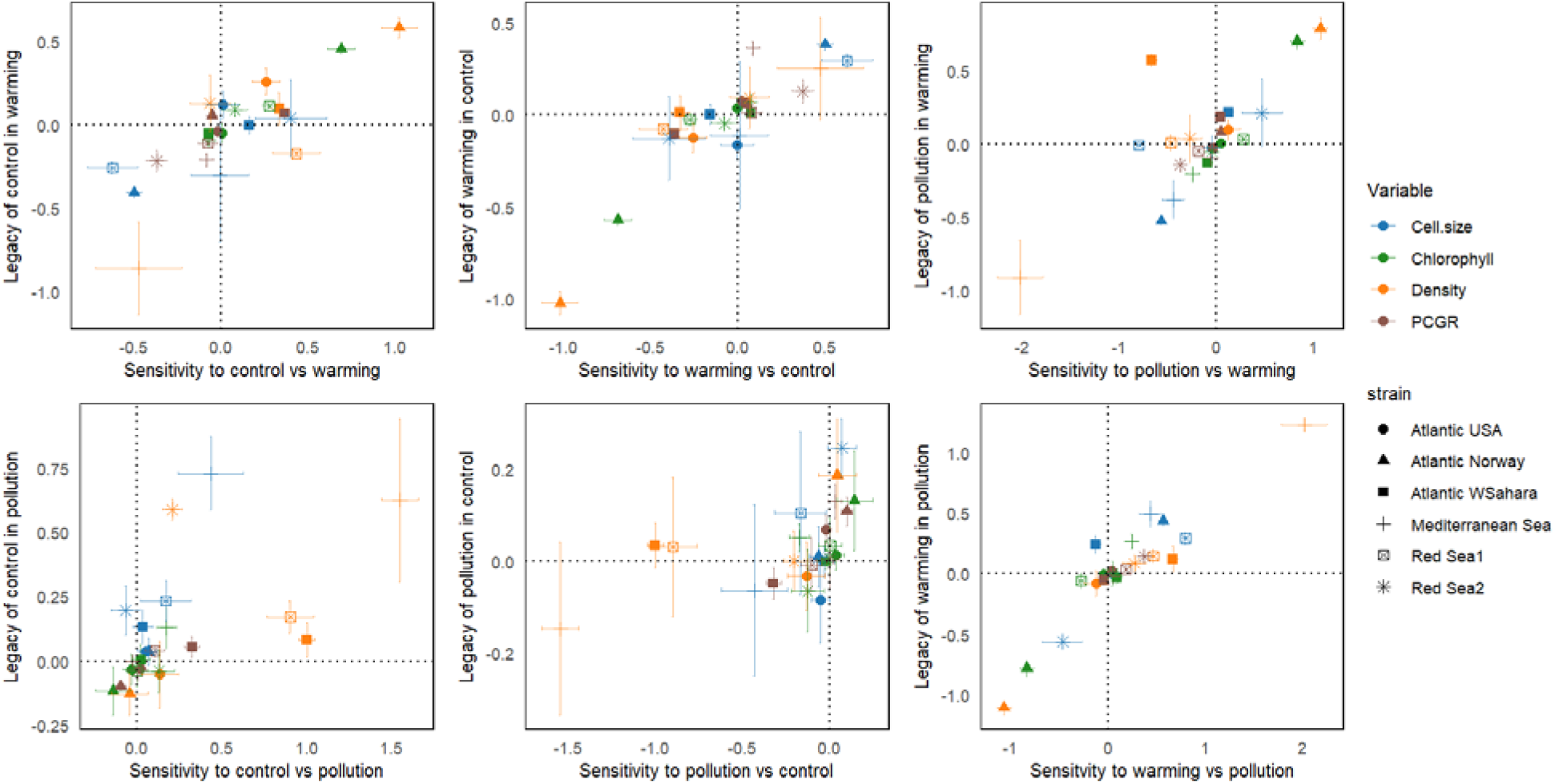
Strain-specific sensitivity and legacy effects in six *Synechococcus* strains. Each panel plots sensitivity (x-axis: difference in response under chronic X vs. chronic Y conditions; XX – YY) against legacy effects (y-axis: difference in response under sequential vs. chronic Y exposure; XY – YY). Points represent mean values (n = 3) ± standard error. Colours indicate response variables (cell size, chlorophyll content, population density, and PCGR); shapes represent strain IDs. Conceptual zones (I–IV) are as defined in Fig1.

Chlorophyll and density showed similar responses (zones I or IV) whereas cell size often showed a contrasting response (Fig 5; TC, PT, TP, CT sequences). These findings suggest that the mechanisms of legacy effects differ between functional traits and growth, and are shaped by both strain- and sequence-specific factors. Overall, our results demonstrate that legacy effects closely mirror environmental sensitivity patterns.

## 4. Discussion

This study investigated the existence and mechanisms of legacy effects during sequential exposures in six Synechococcus populations. Results confirm that legacy effects of past environments significantly alter population growth and trait expression in subsequent environments. Although legacy effects consistently impacted both growth and traits, these responses did not covary, providing conditional support for hypothesis H1. Consistent with hypothesis H2, legacy effects were related to strains’ sensitivity to environmental conditions and could be classified into four distinct patterns, as illustrated in Fig. 1.

### 4.1. Strain-specific sensitivity reflects niche-widths and constrains beneficial acclimation

The six strains in this study originated from five distinct habitats (Table 1), providing a strong basis for examining ecologically meaningful contrasts. Under chronic warming, the Atlantic strains experienced a significant decrease in growth, whereas the Mediterranean and Red Sea strains were largely unaffected (Fig 3). These patterns likely reflect intraspecific variation in thermal niche breadths, shaped by habitat-specific thermal regimes (Krinos et al., 2025; Thomas et al. 2012; Magnuson et al., 1979). Thermal niche theory has been widely used to understand and predict species responses to thermal variability (Gvoždík, 2018). Kremer et al. (2017) showed that phytoplankton share a common temperature sensitivity but differ in their maximum growth rates and thermal niche widths. By extension, *Synechococcus* thermal widths in our study may be similarly broad yet simply shifted: a +2 °C increase moves warmDhabitat strains closer to their T □ □ □, thereby enhancing growth, whereas it pushes coldDhabitat strains past their T □ □ □, reducing performance.

Notably, our results show that prior exposure to a given environment does not always confer a growth advantage when subsequently exposed to the same environment, contrary to the predictions of the Beneficial Acclimation Hypothesis (BAH; Leroi et al., 1994; Wilson & Franklin, 2002). This was particularly evident under chronic pollution and, to a lesser extent, chronic warming (Fig. 3). Chronic exposure to pollution led to markedly negative sensitivity in most strains, except for the Atlantic USA strain (Fig 3), which may have developed tolerance due to potential exposure to atrazine, which is a widely used herbicide in this region (Yang & Zhang 2020; Zhang et al., 2022). According to the BAH, organisms acclimated to an environment should perform better in that same environment than those previously exposed to a different one. In our study, however, performance under chronic conditions was in some cases reduced, which we interpret as a case of negative sensitivity, mediated by exposure type and strain identity. This suggests that the benefits of acclimation are not universal, and may be constrained by underlying physiological limitations as a result of negative sensitivity.

### 4.2. Changes of functional traits and population growth are decoupled across exposure types

Our findings support hypothesis H1: past environments influence both population growth and traits. However, this influence is more complex than our simplistic expectations of a linear increase or decrease. We observed a decoupling of trait and growth responses across chronic and sequential exposures. For example, under chronic warming, Atlantic strains (USA and Norway) showed declines in PCGR and chlorophyll content, while cell size either increased or remained unchanged-breaking the typical expectation that smaller, faster-growing cells trade off pigment allocation (Fig 3). Growth and adaptability trade-offs have been reported in *Escherichia coli* when the composition of growing culture was abruptly shifted (Basan et al., 2021). Chronic pollution, by contrast, led to concurrent reductions in chlorophyll content, cell size, and growth, suggesting that some environments can reinforce train-performance correlations.

Interestingly, under sequential exposures, the timing of trait responses varied (Fig 4). Chlorophyll remained comparative stable regardless of past exposure, while cell size lagged in matching with the subsequent environment. This suggests that pigment synthesis can respond more rapidly to new conditions, while cell size reflects a delayed response. Such trait-specific lag indicates that the temporal dynamics of individual traits are influenced by the exposure type. Emerging evidence for two-way interactions (Holmes et al., 2024), including trade-offs between population growth and traits related to resource allocation (Reimers et al., 2017) support this view.

### 4.3. Legacy effects in sequential environments: beyond beneficial and detrimental outcomes

Our findings support hypothesis 2: for most strains and environmental sequences, legacy effects, defined as performance differences due to a past exposure, aligned with sensitivity. That is, populations that had a strong (positive or negative) sensitivity to an environment typically had the same legacy effect in subsequent environment (Fig. 5).

**Table 3:** Replication statement.

| Scale of inference | Scale at which the factor of interest is applied | Number of replicates at the appropriate scale |
| --- | --- | --- |
| Strains | Strains ($n = 6$) | 162 experimental units - 6 strains X 3 past environments X 3 subsequent environments X 3 replicates. |

**Table 4:** Results of mixed-effects ANOVA testing the effects of strain, exposure type, and their interaction on four response variables: density, cell size, PCGR, and chlorophyll content, under chronic warming (TT), pollution (PP), and control (CC; reference level) conditions. Reported statistics includ numerator and denominator degrees of freedom (df), mean square error (MSE), F-values, generalized eta-squared (ges), and associated *p*-values. Post hoc pairwise contrasts by strain are presented in Supplementary Material Table 1.

| Mixed-effect ANOVA results |  |  |  |  |  |  |  |
| --- | --- | --- | --- | --- | --- | --- | --- |
| Variability due to Chronic exposures compared to control |  |  |  |  |  |  |  |
| Response Variable | Effect | df (num) | df (den) | Mean Sq. Error | F-value | Generalized Eta Squared | p-value |
| Density | strain | 5.00 | 10.00 | 0.01 | 54.99 | .898 | <.001 |
| Density | exposure | 1.98 | 19.75 | 0.01 | 110.82 | .883 | <.001 |
| Density | strain:exposure | 9.88 | 19.75 | 0.01 | 54.03 | .948 | <.001 |
| Cell.size | strain | 5.00 | 10.00 | 0.03 | 67.63 | .935 | <.001 |
| Cell.size | exposure | 1.83 | 18.26 | 0.02 | 9.40 | .351 | .002 |
| Cell.size | strain:exposure | 9.13 | 18.26 | 0.02 | 10.57 | .753 | <.001 |
| PCGR | strain | 5.00 | 10.00 | 0.00 | 45.26 | .866 | <.001 |
| PCGR | exposure | 1.43 | 14.32 | 0.00 | 16.95 | .548 | <.001 |
| PCGR | strain:exposure | 7.16 | 14.32 | 0.00 | 18.40 | .868 | <.001 |
| Chlorophyll | strain | 5.00 | 10.00 | 0.01 | 196.53 | .982 | <.001 |
| Chlorophyll | exposure | 2.00 | 19.98 | 0.00 | 24.84 | .527 | <.001 |
| Chlorophyll | strain:exposure | 9.99 | 19.98 | 0.00 | 31.72 | .877 | <.001 |

Other results revealed that exposure to one environment could enhance tolerance to a different subsequent environment, through amplification and overcompensation. For instance, strains previously exposed to warming performed better under pollution (TP vs. CP), and vice versa (PT vs. CT). This could be a form of “stress-memory” or “phenotypic memory” i.e. lasting effects of stress events on population response (Anderson et al., 2025; Zhou & Wang, 2023). Populations potentially develop shared or interacting response pathways when exposed to different stressors. In case of sequential exposure to different, this is known as “trans-memory effect” (Zhou & Wang, 2023). For co-occurring stressful environments, this is known as cross tolerance and has been widely documented across plants (Kamanga et al., 2022), animals (Rosenberg et al., 2021), and insects (Denny & Dowd, 2022). This outcome is advantageous, as it allows populations to better tolerate contrasting environments in a sequence, a common feature of natural ecosystems (Zhou & Wang., 2023). Recent studies highlight this as a potential pre-adaptive mechanism to global environmental change (Rodgers & Gomez Isaza, 2021; 2023).

Conversely, certain legacies led to reduced performance in sequential exposures relative to chronic counterparts. This was especially apparent in pollution or temperature-to-control sequences (PC, TC). A likely explanation is stress accumulation within cells. Exposure to stressful conditions (in this case warming or pollution), can lead to the buildup of secondary metabolites that slow down growth as a protective mechanism (Yadav et al., 2022). Additionally, prior exposure to warming in some strains may cause metabolic shifts via membrane fluidity and lipid metabolism as a defence system (Lauritano et al., 2020). Stressed populations may have activated these protective mechanisms, but then been unable to rapidly turn them off when the stress ceased, resulting in continued lower growth. Similar findings have been reported in *Drosophila nepalensis* (Ramniwas et al., 2022).

Collectively, our findings highlight that legacy effects are mechanistically diverse and context-dependent. Population responses during sequential exposures can reflect amplification, overcompensation, constraint, or depression, contingent on environment(s), sequence order, and strain origin. This complexity underscores the need to incorporate legacy effects and their underlying mechanisms into predictive models of organismal responses under increasing environmental variability.

## Conclusions

In this study, we examined how legacy effects shape population responses during sequential environmental change. Our findings demonstrate that legacy effects significantly influence population growth and functioning with outcomes linked to strain-specific sensitivity which may be rooted in its habitat of origin. We speculate that *Synechococcus* strains from warmer oceans, such as the Mediterranean Sea and Red Sea, may better withstand chronic or sequential warming than those from the Atlantic Oceans. Strains from the USA coast of Atlantic may maintain growth in polluted environments compared to strains from Mediterranean or European coasts of Atlantic.

Future work should incorporate additional functional traits (beyond growth, chlorophyll content, and cell size) relevant to *Synechococcus* sp. ecophysiology (e.g. photosystem efficiency, stress protein expression) and explore molecular mechanisms such as methylation patterns that may underlie strain-specific trait plasticity. Integrating bioassay-based sensitivity data with legacy-effect models offers a promising path forward for forecasting population persistence under real-world environmental variability.

As climate change intensifies environmental variability, understanding population responses in dynamic, sequence-dependent contexts is essential. Our results underscore the need to move beyond single-stressor approaches and toward sensitivity-based models that capture legacy effects such as overcompensation, amplification, and depression to improve predictions of population trajectories.

## Notes

### Competing Interest Statement

The authors have declared no competing interest.

