## Supplementary figure 1. 2 for "Trait and Growth Responses to Sequential Environmental Change linked to Sensitivity in *Synechococcus* populations"

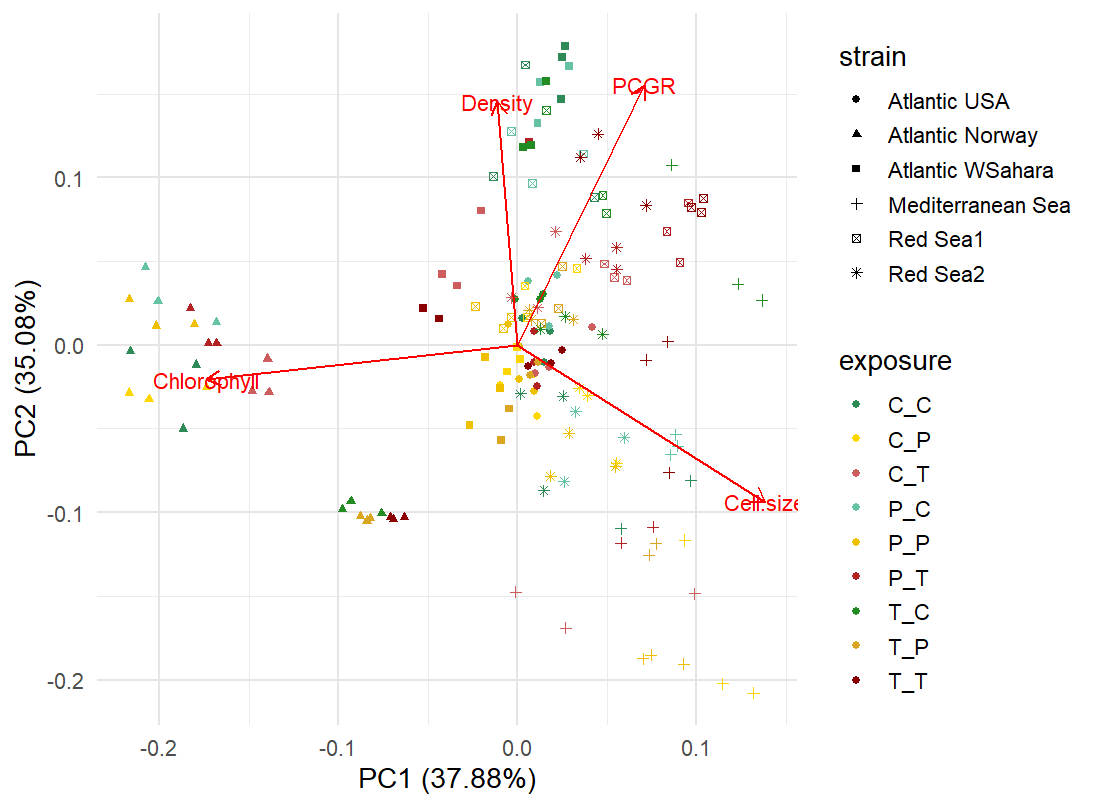

**Fig 1: PCA of four variables for this study, colored by exposure type and shaped by strain ID.** PC1 and PC2 explain ~73% of the total variance. Shapes represent habitat origin and colors indicate sequence categories. Red vectors show the contribution of each variable (e.g., density, pcgr, cell size and chlorophyll) to sample variation. Clustering patterns suggest strain-specific structuring and overlapping exposure-type distributions.

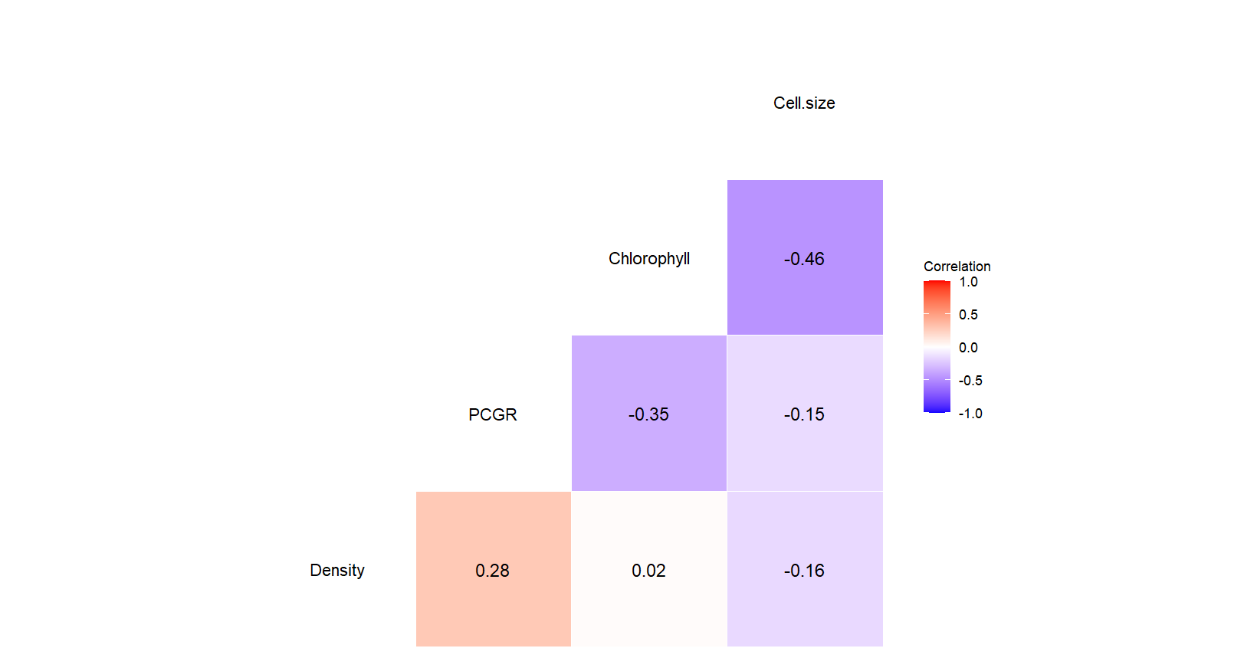

**Fig 2: Pairwise correlation matrix of response variables across exposure type and strain IDs.**
Pearson correlation coefficients (r) are shown for per-capita growth rate (PCGR), population density (density), chlorophyll content (chlorophyll), and cell size, calculated on the full set of strainID-exposure type combinations. Color shading indicates the strength and direction of each correlation (blue = strong negative; white = zero; red = strong positive), with exact r-values overlaid (rounded to two decimal places).
